# Inhibitory neurons are a Central Controlling regulator in the effective cortical microconnectome

**DOI:** 10.1101/2020.02.18.954016

**Authors:** Motoki Kajiwara, Ritsuki Nomura, Felix Goetze, Tatsuya Akutsu, Masanori Shimono

## Abstract

The brain is a network system in which excitatory and inhibitory neurons keep the activity balanced in the highly non-uniform connectivity pattern of the microconnectome. It is well known that the relative percentage of inhibitory neurons is much smaller than excitatory neurons. So, in general, how the inhibitory neurons can keep the balance with the surrounding excitatory neurons is an important question.

We observed effective networks, reflecting causal interactions, of ~1000 neurons in cortical acute slices. Surprisingly, we found that inhibitory neurons are not only located at more central positions than excitatory neurons but also have stronger controlling ability of other neurons than excitatory neurons. Besides, we found that the precedence in centrality and controlling ability of inhibitory neurons are well observed in deep cortical layers by comparing with distribution of neurons coloured by NeuN immunostaining data. Preceding the observation, we also found that inhibitory neurons show higher firing rate than excitatory neurons, and that their firing rate also closely obey a log-normal distribution as previously known about excitatory neurons. Additionally, their connectivity strengths also obeyed a log-normal distribution.

In summary, within the network interaction of huge numbers of neurons, inhibitory neurons seem to produce a central controlling system that sustains the homeostatic behavior of the brain. A similar evaluation in different life stages and in disease states *etc.* will not only provide deeper understandings in the homeostasis of the brain, but also will provide a selective and effective way to stimulate individual neurons to modulate neuropsychiatry or neurodegeneration disease states.

## 1. Introduction

### 1-1. Initial statement

The brain is a highly non-random network system. Inside of the system, electrical information flows on the networks in a highly balanced manner under mutual interactions between the excitatory and inhibitory neuron pools. Therefore, the extraction of the rules that exist behind the distributions of these two types of neurons within the functional network organization is a crucial scientific topic.

### 1-2. E/I balance and disease

From over two decades ago, E/I (excitatory and inhibitory) balance has been regarded as the key factor not only for epilepsy [Sloviter, 1987], but also for various mental diseases such as schizophrenia [Lewis et al, 2005] and autism [Rubenstein, Mersenich, 2003; Nelson, Valakh, 2015]. Beyond the phenomenological observations, causal influences from the E/I imbalance at the medial prefrontal cortex to behavioral deficits were also recently experimentally verified [Yizhar et al., 2011]. Historically, it had been known that suppression of inhibitory neurons causes less selectivity for external stimuli [Sillito, 1975]. However, it is also true that E/I balance is an over-simplified quantification when considering the inherently wide diversity and non-uniformity of huge numbers of neurons [Nelson, Valakh, 2015; O’Donnell et al., 2017]. As a fact, relative positions of individual neurons are highly non-uniform, and rare outstandingly influential neurons sometimes have exceptional ability to influence the global brain dynamics [Miles, Wang, 1983]. The potentially related widely-spreading structural projection pattern was also recently demonstrated [Reardon, 2017]. Therefore, the global mean will not easily represent such system properties.

### 1-3. Network theoretical approach

To quantify such highly non-uniform architectures, network theory has been widely applied to brain data, especially at macroscopic scale [Bullmore, Sporns, 2009; Kaiser, Hilgetag 2006] and also more widely beyond neuroscience [Newman, Barabasi, Watts, 2006]. When we zoom into the microcircuits of neurons, several previous studies demonstrated complex intertwined architectures and have also extracted comprehensive designs and their rules [Song et al., 2005; Shimono, Beggs, 2011, 2014]. Previous studies of functional connectivity patterns demonstrated the small-worldness, namely the co-existence of short path-length and high clustering property in the cortex [Yu et al., 2008], scale-free like topology and existence of hub neurons in the hippocampus [Bonifazi, et al., 2009], log-normal distribution [Buzsaki, Mizusaki, 2014], and also structure-function relationship among convergent connections [Bock et al., 2011]. In summary, these past studies of local neuronal circuits have discovered various specific non-uniformity of neuronal networks. However, our knowledge about the difference of contributions between E/I neurons in terms of relative positions in a large population of neurons is yet fairly limited. Specifically, there are few studies about effective networks as we will mention later.

### 1-4. Data acquisition and analysis scheme

This research evaluated how E/I neurons are located differently in the effective networks of neurons by applying several graph theoretical analyses. This analysis scheme allows us to quantify different roles of individual neurons in terms of the relative positions and interactions with the many other neurons. The effective network was reconstructed from neuronal spikes recorded by a cutting-edge Multi-Electrode Array system, and we selected TE (Transfer Entropy) to estimate the effective networks, quantifying the directed causal coupling strengths, among neurons. The connectivities defined from neuronal activity can be categorized into functional connectivity and effective connectivity. Among them, the functional connectivity simply evaluates statistical dependence of dynamic couplings using correlation, covariance, spectral coherence, or phase lockings between two time series. Although they can change even on the short time scales, the evaluation does not explicitly quantify the causal influences. Effective connectivity explicitly deals with the causal influences between more than two time series [Friston, 1994; McIntosh, 1991; Aertsen, 1989]. TE has several preferred abilities for the estimation ability [Lungarella et al., 2007], and for theoretical analyses [Verdes, 2005; Vicente, 2011; Lizer et al., 2008], and TE has been widely adopted in various neuroscientific studies. Although there are many strengths of TE, it cannot distinguish between excitatory and inhibitory connections. The current study applies a new analysis scheme [Goetz, Lai, 2019] to our experimental recordings of neuronal spike data.

### 1-5. Main questions

Certainly, the estimation of connectivity will not be the final destination of neuroscience. The main purpose of this study is to explore how differences of relative positions within the networks of excitatory and inhibitory neurons play different roles in microconnectomes, the comprehensive connectivity patterns of neurons. Evaluations of connectivity patterns provide more information than simple differences of their spatial adjacency, such as participating layers.

When comparing between excitatory and inhibitory neurons in the cortex, it is well known that the number of excitatory neurons is much larger than inhibitory neurons. In such an environment, how can inhibitory neurons keep the balance with excitatory neurons?

It is often discussed that inhibitory neurons have higher firing rate than excitatory neurons to keep the E/I balance, however, it is not yet well-known how the relative position of inhibitory neurons helps to keep the E/I balance.

This study will answer this question by passing through the following three steps: First, we evaluate the basic statistical properties of neuronal ensembles. The most fundamental property of neuronal activity would be firing rate, and the most fundamental property of network architecture would be the weight distribution and the degree distribution.

Our past studies demonstrated that neuronal firing rates and connectivity weights follow a log-normal distribution, and that the degree distribution shows long-tailed form. We specifically test if these properties sustain or not when we observe excitatory and inhibitory neurons separately.

Second, we quantified the differences of relative positions of excitatory and inhibitory neurons (E/I neurons) within the comprehensive network architecture of the microconnectome. Especially because we are interested in the relative difference of “influence” of E/I neurons to other neurons, how central they are evaluated based on K-Core Centrality (KC).

Third, we evaluated how much E/I neurons are relatively more influential in terms of their ability to drive or control other nodes in the network system based on the Feedback Vertex Set (FVS) method. Note that FVS is a well-known concept in graph theory [Slater, 1961; Karp 1972], and its relations with attractors and controllability to attractors were shown in [Akutsu et al., 1998] and in [Mochizuki, et al., 2013], respectively. Furthermore, it was demonstrated that genes and proteins corresponding to FVS nodes tend to play important biological roles [Mochizuki et al., 2013; Bao et al., 2018; Zanudo et al., 2017].

From these steps, we could reveal that inhibitory neurons are not only more active than excitatory neurons but also that they are located in a more central position, and also that they have higher controlling ability of the entire network dynamics in local cortical neuronal circuits.

## 2. Materials and Methods

### 2-1. Animals

We used female C57BL/6J mice (n=7, aged 3-5 weeks). All animal procedures were conducted in accordance with the guidelines of animal experiments of Kyoto University, and have been approved by the KU Animal Committee [fig. 1-a].

**Figure 1.**
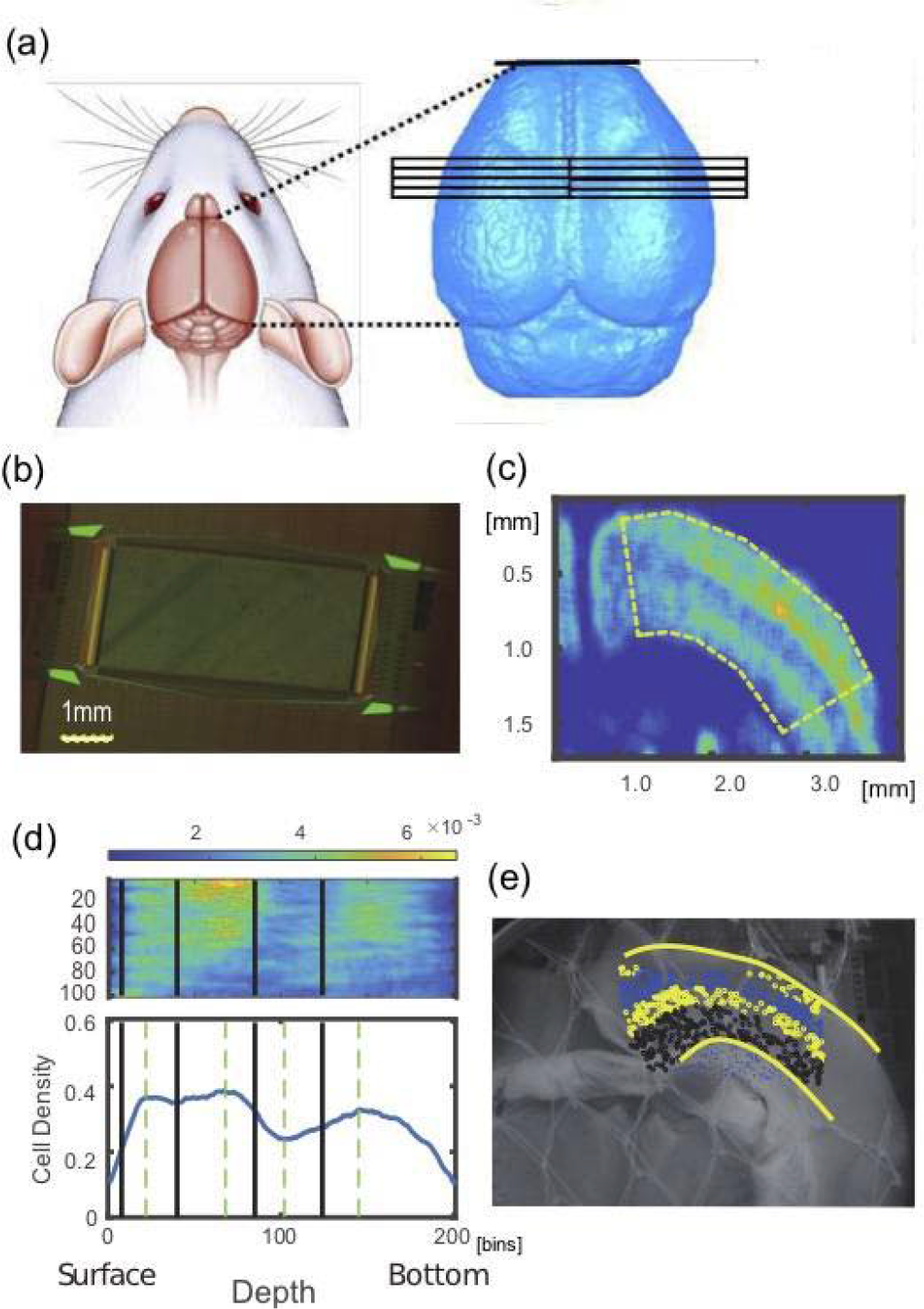
Data samples, slices, staining and layer detections: (a) shows anatomical coordinates where we prepared slices from. The left and right hemispheres were equally selected. (b) is a photo of a Multi-electrode array used in this study. (c) is a smoothed NeuN staining image. We can find the non-uniformity of density of neurons. The curved regions as shown dotted line in the panel (c) was re-shaped into a straight box as shown in the top panel in (d), and the density was expressed as sum on the vertical axis against layer boundaries in the lower panel of (d). Then, we determined the 4 layer boundaries shown as thick lines in the lower panels, and mapped onto the non-stained photo image (e) and all neurons identified from spike sorting process were categorized into individual layers.

### 2-2. Induction of anesthesia

We anesthetized mice with 2% isoflurane in air flowing through the box at a flow rate of 1.4 L/min using an anesthesia system, which is composed of an isoflurane vaporizer, air flow meter, and air compressor. Then, mice were located in an acrylic anesthesia box attached on a heater mater adjusted to 37 °C. After checking the anesthesia is sufficient, we placed the mouse in an MRI-compatible cradle in the prone position.

### 2-3. MRI acquisition

MR experiments were conducted on a 7 T, 210 mm horizontal bore, preclinical scanner (BioSpec 70/20 USR, Bruker BioSpin MRI GmbH, Ettlingen, Germany), equipped with a gradient system of 440 mT/m at 100 μs ramp time. A quadrature volume resonator (inner diameter 35 mm, T9988; Bruker BioSpin) was used for RF excitation and signal reception. MRI data was acquired with ParaVision 5.1 software (Bruker BioSpin). Three dimensional T_2_ weighted (T2W) images of the mouse whole brain were acquired with relaxation enhancement (RARE) sequence [fig. 1-a]. The acquisition parameters were as follows (based on the protocol TurboRARE-3D; Bruker BioSpin): repetition time (TR), 2000 ms; echo time (TE), 9 ms; effective TE, 45 ms; RARE factor, 16; acquisition matrix size, 196 × 144 × 144; field of view (FOV), 19.6×14.4×14.4 mm^3^; acquisition bandwidth, 75 kHz; axial orientation (coronal orientation in scanner setting): fat suppression with 2.6 ms-gaussian-shaped π/2 pulse with 1051 Hz bandwidth followed by spoiler gradient; 2 dummy scans; number of averages, 3; acquisition time, 2 h 42 m. 2.59-ms excitation and 1.94-ms refocusing pulses were used. The shape of their pulses was sinc-3 shape multiplied by a gauss function with a 25% truncation level (sinc3) and their bandwidth was 2400 Hz.

### 2-4. Preparation of experimental solutions

Preceding the neuronal activity recordings, we prepared two solutions, an ice-cold cutting solution (CS) and an artificial cerebrospinal fluid (ACSF). The necessary ice-cold CS was 500 mL for each experiment, and it was a mixture of the following chemical materials; 2.5 mM KCl, 1.25 mM NaH2PO4, 7 mM MgCl2, 15 mM glucose, 25 mM NaHCO3, 0.5 mM CaCl2, 11.6 mM sodium ascorbate, 3.1 mM sodium pyruvate, and 100 mM choline choline chloride, bubbled with 95% O2 and CO2. The pH was adjusted to 7.3-7.4. We also prepared 1 L ACSF, and the content is 127 mM NaCl, 2.5 mM KCl, 1.25 mM NaH_2_PO_4_, 1 mM MgCl_2_, 15 mM glucose, 25 mM NaHCO_3_, and 2 mM CaCl_2_, bubbled with 95% and 5% CO_2_. ACSF was also warmed (~34 °C) and adjusted the pH to 7.3-7.4.

### 2-5. Slice preparation and Spike recording

To start neuronal spike recording using a MEA system, we extracted brain after sufficiently anesthetizing with 1%-1.5% isoflurane, and then transferring it to a Petri dish (100 mm × 20 mm) filled with the ice-cold cutting solution in which air containing 95% O_2_ and 5% CO_2_ flowing. We made 2-5 coronal slices (300 μm thick) from the extracted brain using a vibratome (Neo Linear Slicer NLS-MT; DOSAKA EM CO., LTD). To optimize the cutting speed, frequency, and vibration amplitude of the system we set them as 12.7 mm/min for the speed, 87-88 Hz for the frequency, and 0.8-1.0 min for the swing width, respectively. When we temporarily needed to cut a brain carefully, we additionally slowed down the cutting speed. While cutting the brain slices, we recorded the sliced coordinates in the format, including the anterior-posterior coordinates, hemisphere, and other conditions. All slices, analyzed in this study, were taken from the dorsal cortex including the barrel sensory area and the primary motor area [fig. 1-a]. Then, we gently transferred the slice to a beaker filled with prewarmed (~34 °C) ACSF using a thick plastic pipette, and incubated them in the beaker at ~34 °C for 1 h.

After 1 h has passed, we moved a slice on the chip using a thick plastic pipette, and set the position using a soft brush to properly record from specific brain regions including cortex. The outline of the array was rectangular (2.0mm×4.0mm) and 26000 electrodes were uniformly arranged [fig. 1-b], and the inter-electrode distance was 15 μm (Maxwell Biosystem, MaxOne) [Obien et al., 2015]. Preceding the main recording, we performed so called “pre-scan” as a combination of 30[sec] recordings from densely distribution 1020 channels included in 25 groups, and selected sensors showing stronger responses than 0.1 Hz and 0.02 mV for the main recording. Then, we started the main recording of spontaneous neuronal activities from selected up to 1020 sensors for ~2.5 hours. During recording, the slices were perfused at 1mL/min with ACSF that was saturated with 95% O_2_/5% CO_2_ while controlling the temperature of the perfusate around 34°C.

### 2-6. Brain surface scan

We recorded a 3D scan surface at three times using a 3D structured light technology scanning system (SCAN in a BOX; Open Technologies). We performed 3D scan recording at three times; The three recordings are whole brain, brain blocks after cutting into two blocks and before making slices, and brain blocks that remained after making multiple brain slices. The whole process was performed based on the 3D novel embedding overlapping (3D-NEO) protocol [Ide al., 2018]; The first 3D scan recording was done just after extracting a whole brain from a mouse head. Here, we removed the brain and gently dropped it into a Petri dish with ice-cold cutting solution (CS) after decapitating from a mice fully anesthetized with 1.0−1.5% isoflurane. The second recording was performed from two brain blocks cut in the middle of the coronal plane. The cut brain blocks were attached on the surface of brain-block-base (BBB), a 6.6 cm^2^ square and 0.5 cm^2^ thick magnet ring with sticky tape, using instant glue. We prepared such base to transfer the brain blocks smoothly from 3D scanning to the brain slicer [Ide al., 2018]. Third 3D scan recording was performed from the remaining brain block after all slicing processes. Unexpectedly, the last one is the most important for knowing the position of the last slice, and accordingly positions of other 2-4 slices. Before scanning, we gently wiped fluid from the brain surface using the microfiber cloth, which is important to effectively absorb water which causes diffuse reflections. In all scans, the rotating angle was 22.5 degrees, and we performed an automatic coregistration among images taken from 16 different angles. After scanning, we checked that all scanned surfaces are nicely overlapped with each other.

We processed the scanned images using the 3D scan and processing software IDEA (including SCAN in a BOX). First, we integrated small mismatches among 16 image using the automatic alignment option by IDEA. Then, we merged the objects of integrated images recorded from above and below or frontal and occipital, using the manual alignment option. The optimization algorithm is the iterative closest point (ICP) algorithm without nonlinear deformation as frequently applied to hard 3D objects [Besl, McKay, 1992]. Finally, we made a meshed object with high resolution from the merged object by the mesh generating option, and saved it as stl binary format to compare with MRIs for individual brains.

### 2-7. Spike Sorting

We performed offline spike sorting using the SpyKING CIRCUS software. The main strength of the algorithm utilized in the software is high accuracy even though the calculation cost is low by categorizing spikes into neurons based on a template-matching method, in which templates were produced as representative waveforms averaged within a part of time series [Yger et al., 2018]. The low computational cost was beneficial for our large number of electrodes (>1000).

First, spikes were detected as threshold crossings of a 6 standard deviation line from the high-pass filtered time series with a Butterworth filter and also whitening to delete pseudo correlation originated from external noise using 20 second data where spikes did not appear. The order of the filter was three and cutoff frequency was 500Hz. Then, we isolated waveforms as 5 time sections around the randomly chosen peak points. Second, we performed clustering for individual groups of waveforms which were once divided into several groups located within a 150 [*μm*] radius This step was implemented in Spyking CIRCUS to reduce necessary computational memory. The waveform data, maximally 10000 samples, were projected into five dimensional principal component space. Third, we performed a density-based clustering [Yger et al., 2018], which gathers other samples surrounding the peaks after estimating local density peaks locating at centers of individual clusters [Rodriguez, Laio, 2014]. We also iteratively merged different clusters, in which “normalized distance” is smaller than 3. Fourth, we estimated templates as two dimensional descriptions of individual clusters. The two dimensions consist of average extracellular waveform and the variance. The second component is orthogonal to the first component in the sense of the, so called, Schmidt decomposition. Based on similarity with the first component, so the waveform, of the template and the raw data, we seeked individual spikes originated from the same putative neurons one by one. Fifth, we merged two neurons if different templates were too similar (correlation > 0.975). Finally, we eliminated neurons whose firing rate is lower than 0.2Hz because there could be somehow noisy samples.

### 2-8. Immunohistochemistry staining and layer extractions [k,f,n]

#### 2-8-1. Immunohistochemistry staining [n]

On the same day where the MEA recording is finished, we fixed the slices with the 4% paraformaldehyde (PFA) in phosphate-buffered saline (PBS) overnight at 4°C. On the second day, we incubated it with antigen retrieval solution (10 mM sodium citrate, pH 6.0) for 20 min at 95~98°C after washing the slices 3 times with PBS for 5 min in total, and cooled it to the home temperature. Then, we prepared the blocking solution by adding 4% normal goat serum (NGS) and 0.5% octylphenol ethoxylate (Triton X-100) to the 1% Na bisulfite-Tris solution (pH 7.5). After washing the slices with 1% Na bisulfite in 50 mM Tris-buffered saline, we incubated the slices with the blocking solution for 2 hr at room temperature (20~25°C). We also prepared the primary antibody solution by adding 1% NGS and 0.5% Triton X-100 to the 1% Na bisulfite-Tris solution and diluting primary antibodies. Now, we incubated the slices with the primary antibody solution, consisting of 1:800 of mouse anti-GAD67 and 1:500 of rabbit anti-NeuN, overnight at 4°C. Please notice that this study used information only of NeuN staining. On the **third day**, we prepared the Tris-NaCl solution (8.5 g/L NaCl in 50 mM Tris-buffered saline, pH 7.5) and the secondary antibody solution by adding 3% NGS and 0.3% Triton X-100 to Tris-NaCl and diluted secondary antibodies, consisting of 1:500 of goat anti-mouse and 1:500 of goat anti-rabbit. Then, we washed the slices 3 times with the Tris-NaCl solution for 5 min in total and incubated it overnight at 4°C. On the **fourth day**, we washed the slices 4 times with the Tris-NaCl solution for 5 min in total. Then, we mounted the slices on slide, embedded with antifade reagent (SlowFade Gold; invitrogen), and enclosed with a coverslip, and recorded the distribution of cell bodies at a magnification of 10x from the slides using a fluorescence microscope (All-in-One Fluorescence Microscope BZ-X710; Keyence).

#### 2-8-2. Image processing for extracting layer categories [t, n]

We applied images processes consisting with four steps to the given fluorescent images: First, we extracted positions of cell bodies from original NeuN staining images. Second, we defined layers based on distribution densities of excitatory or inhibitory cell bodies. Third, we overlapped the layer divisions from staining data onto photo images taken just after MEA recordings. Fourth, we overlapped the MEA coordinates onto the photo image, and also gave layer categories for all electrically recorded neurons. Let me explain in more detail about the individual steps;

As we mentioned, **the first step** <u>detects cell positions from staining images</u>. The imaging processes consists of the pre-processing to reduce noises, and the main detection processes. The noise reduction eliminates small masses of bright pixels after filling the gaps or holes generated by random noises. Then, we identified cell bodies’ positions using Watershed algorithm. So, we first drew contour lines that gradually rise from the surfaces of cells toward the center of cells. Using the “height” given by the contour lines, we next detected boundaries between cells overlapping each other. Then, we could define centers of cells from the detected cells’ external boundaries or regions.

The following **second step** aims to <u>draw boundaries of layers based on cells’ density distributions</u> from NeuN images. So, we first defined the density distribution by smoothing the distribution of cells as number of cells existing in 50×50 pixel regions. To define boundaries of layers, we once extended the curves corresponding with surface and bottom of the stained image to straight lines. Then, we sum the number of cells in the direction perpendicular to the cortical layer to obtain one curving line of cell density histogram. The cell density histogram typically shows three convex points and two concave points between the convex points commonly for all cortical samples [Fig. 1-c, bottom panel]. The first convex point counted from the cortical surface indicates the center of layer3, and the second convex point locates at the middle of layer 4, and the third convex point far from the cortical surface involved in the layer 6, and two concave points individually corresponds to the boundary between layer 2/3 and layer 4, and the middle point of the layer 5. So, we defined all boundaries as the following steps; First, between the cortical surface and the first convex point, we regarded the 1/3 from the cortical surface as the boundary between layer 1 and layer 2/3. Second, as mentioned, we defined the boundary between layer 2/3 and layer 4 at the concave point between the first and second convex point. Third, we defined the boundary between layer 4 and layer 5 at the middle between the second convex point and the second concave point. Fourth, we defined the boundary between layer 5 and layer 6 at the middle between the second concave point and the third convex point [Fig. 1-c, bottom panel]. The produced boundary lines were mapped on the original staining images [Fig. 1-b].

Because we have a set of boundaries, as **the third step**, we needed <u>to map the boundaries onto photo image taken just after the electrical recording</u> [Fig. 1-d], and also onto the coordinate on the Multi-electrode array [Fig. 1-a]. We simply call the photo taken just after the electrical recording as non-stained image here.

To overlap stained image and non-stained image, we selected common typical points for them (10 points on the cortical surface and 7 points on the bottom of cortex, and other maximally other typical points). By optimally mapping each other based on iterative closest point algorithm [Symon et al., 2001], we got a transformation matrix from the stained image to the non-stained image.

The final **fourth step** is just to <u>overlap non-stained image and MEA coordinates</u>. So, we selected four corners in the non-stained image, and got the transformation matrix from the MEA coordinate to the non-stained image. Then, finally, all neurons estimated from the spike sorting process could be mapped onto the non-staining image, and they met with layer boundary coming from the stained image there [figure 1-d].

### 2-9. Effective connectivity estimation

How to evaluate the causal influences among different variables beyond spurious correlations or confounding factors is a crucial issue [Okatan et al., 2005; Hlavackova, Schindler et al., 2007; Pillow et al., 2008; Gerhard et al., 2011]. We selected transfer entropy (TE) to quantify the causal influences because several previous studies have demonstrated its advantages both in terms of the theoretical formulation and regarding the requirements imposed by the physiological data [Lizier et al., 2008; Vicente et al., 2011; Wibral et al., 2013]. Past studies using Transfer Entropy have successfully demonstrated the structure-function relationship [Garofalo et al., 2009; Stetter et al., 2012; Honey et al., 2007; Ito et al., 2011], consistent topology with patch-clamp experiments and existence of hubs [Shimono, Beggs, 2014], and also a log-normal distribution of connectivity weights [Nigam et al., 2016].

TE is positive if including information about neuron J’s spiking activity improves the prediction of neuron I’s activity beyond the prediction based on neuron I’s past alone (Fig. 2b). The equation for TE used in this study is expressed as:

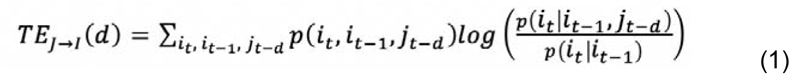

**Figure 2.**
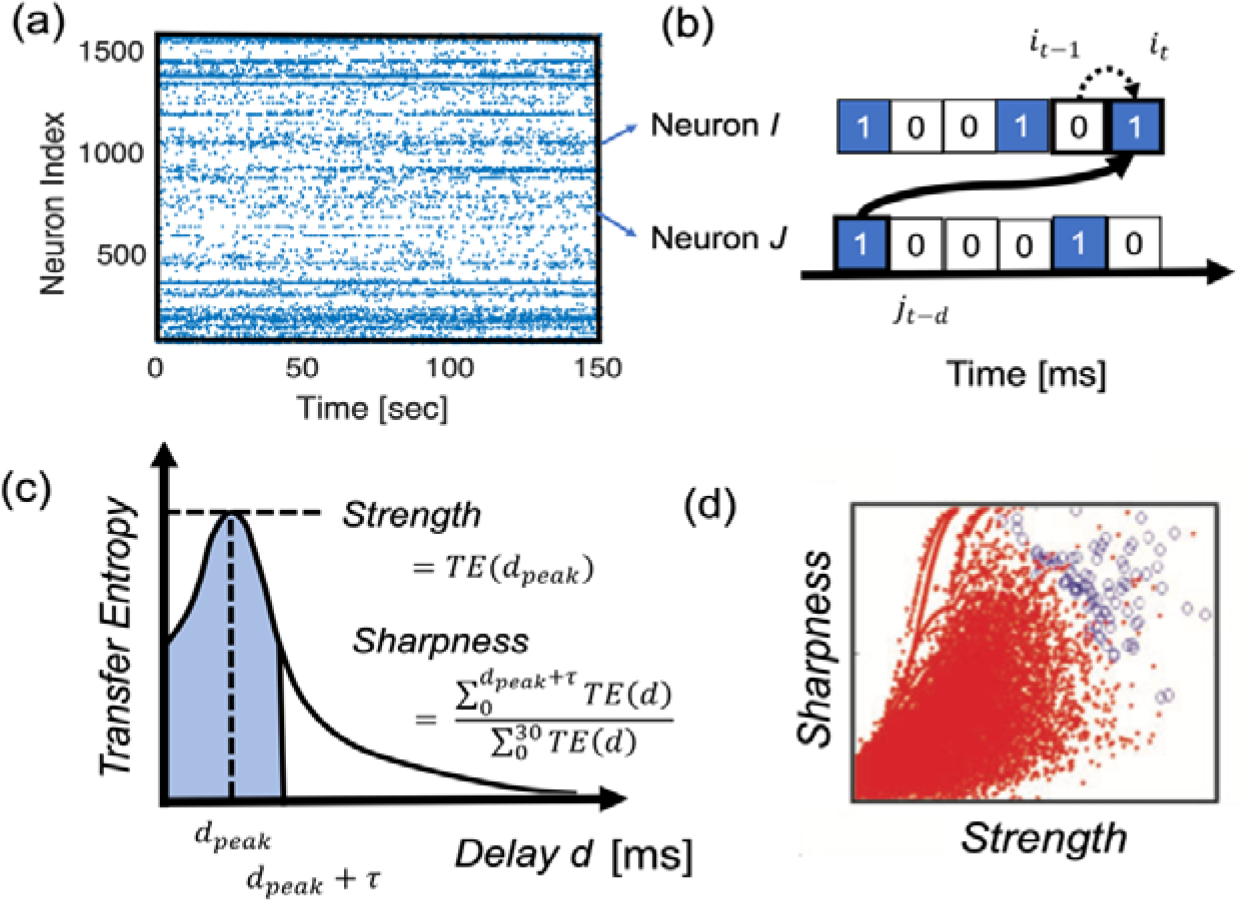
Effective connectivity definition: (a) A raster plot of neuronal spikes given after spike sorting analysis. (b) zoomed-in view of a pair of neurons, I and J. The transfer entropy used in this study specifically quantifies information flow sent from neuron *J* to I that takes delay *d amount of time steps* for the signal transmission, after conditioning on the past activity of neuron *i* at time *t−1*. (c) Computing TE at different delay d, we can get a histogram of Transfer Entropy. To characterize the histogram in terms of strength and sharpness of connections, we characterized two variables shown in this panel. τ is 4 ms in this study. The value was selected as the standard deviation of all TE histograms. (d) The Strength and Sharpness are mapped onto the two dimensional space and we selected connected pairs of neurons shown as blue circles by comparing with the distribution of shuffled spike data (red dots).

This equation quantifies the expected value of the local transfer entropy [Schreiber, 2000; Lizier et al., 2008] 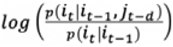 for all, and. The variables *i* or *j* given the index of time t express if the neuron *I* or *J* is active or not at a time bin, and the values are labeled 1 when they are spiking, and 0 when they are not spiking. The time bin sizes are selected as 1ms. The numerator inside of the logarithm expresses the probability of event *i*_*t*_ in neuron *I* conditioned on its own past and the event of the potentially presynaptic neuron J. The denominator expresses the probability of event *i*_*t*_, but conditioned only on its own past event. Schreiber introduced that the difference between these probabilities, as computed by the Kullback-Leibler divergence, which he called Transfer Entropy (1) [Schreiber, 2000], gives a quantity that estimates the information transfer from the source variable *J* to the target variable *I*. (Fig. 2b). Given that synaptic delays between cortical neurons could span several milliseconds [Bartho et al., 2004; Isomura et al., 2006], we adopted a version of TE that allows delays between neuron *I* and *J*.

In the general case, we can include older past events of *i* and *j* in the conditional term as given in the original definition of TE [Schreiber, 2000]. This study used only one time step in the past of neuron *I* and one time step at time delay *d* from neuron *J*. This is because our past study demonstrated that the connectivity matrix produced from higher-order TE, which includes more past events, was very similar to that produced by first order TE, which includes only one time step [Shimono, Beggs, 2014; Nigam et al., 2016].

Here the other terms were as defined in Equation (1). The histogram of TE between 2 neurons was plotted as a function of different delays *d*, which often showed a distinct peak (Fig. 2c). Please note that this delay is essential to determine the directionality of TE because it comes from the fact that the past of neuron J is used to predict the future of neuron I. Thus, connections will be directed from past to future. Besides, how strong and sharp the peak in the histogram is, becomes essential when determining if a direct causal influence through a synaptic connection exists from neuron J to neuron I or not. Therefore, we characterized the *Strength* as the peak value of TE, and the *Sharpness* with the following equation [Fig. 2-(b)]:

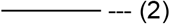

The Sharpness was previously called Coincidence index, however in this paper we use the term *Sharpness* in order to express the intuitive meaning more clearly (fig. 2-(c)). The time window for seeking a peak was 30 ms as the estimated time window for transmitting electrical signal in the spatial size of our recording brain regions [Mason et al., 1991; Swadlow, 1994]. We also used a value of τ = 4 ms, which is the standard deviation for all pairs of neurons. Our past studies selected connections having a strong and sharp peak in the TE histogram on the *Strength-Sharpness* space by comparing the raw connectivity matrix with connectivity matrices generated from shuffled data of spike sequences of presynaptic neurons within ±10ms (fig. 2-(d)) [Shimono, Beggs, 2014; Nigam et al. 2016; Timme et al., 2016]. The jittered data gives the baseline to define the bias-adjusted strength of the connection more accurately according to 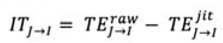. We simply call this value as strength in the following manuscript. These connectivities and strengths have been repeatedly evaluated by comparing with EPSPs measured in patch-clamp recordings [Shimono, Beggs, 2014, Nigam et al., 2016]. We extend the analyses further in order to also apply to inhibitory neurons in the following two subsections.

### 2-10. Positive/negative sign estimation

Extending on past studies, we additionally calculated a new quantity, the sorted local transfer entropy (SLTE). The essence of SLTE is that it sorts the local transfer entropies [Lizier et al., 2008] according to their sign for different interactions. Local transfer entropy estimates the information transfer between two events, rather than two variables, so it can quantify how informative the event of a presynaptic spike is to the event of a postsynaptic spike. For excitatory interactions this yields a positive local transfer entropy, whereas for an inhibitory interaction this would yield a negative local transfer entropy. SLTE then specifically distinguishes between inhibitory influences from excitatory influences, by using a sorting method that takes into account the reversed signs of the local transfer entropies for the excitatory and inhibitory interactions [Goetz, Lai, 2019].

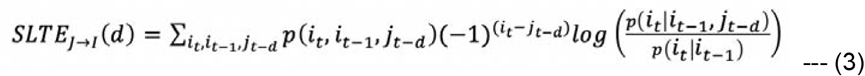

SLTE is similar to the normal Transfer Entropy except that the local transfer entropies are weighted differently, according to the multiplier. For our case where *i*_*t*_ and *j*_*t−d*_ are 1 or 0, depending on whether the respective neuron’s event is spiking or inactive, this multiplier becomes +1 when both *i*_*t*_ and *j*_*t−d*_ have the same event and −1 when the events are different. This sorting rule then makes all terms in the sum of SLTE positive for excitatory interactions and negative for inhibitory interactions, where observing the same events (e.g. a postsynaptic spike at *i*_*t*_=1 that follows a presynaptic spike at *j*_*t−d*_=1) yields a positive local transfer and observing unequal events (e.g. a presynaptic spike at *j*_*t−d*_=1 not resulting in a postsynaptic spike at *i*_*t*_=0) yields a negative local transfer entropy for excitatory interactions. Similarly, for inhibitory interactions the situation is reversed, so that all terms in the sum of SLTE become negative with the sorting rule. Here, observing the same events (e.g. a postsynaptic spike at *i*_*t*_=1 that follows a presynaptic spike at *j*_*t−d*_=1) yields a negative local transfer entropy, whereas observing unequal events (e.g. a presynaptic spike at *j*_*t−d*_=1 not resulting in a postsynaptic spike at *i*_*t*_=0) yields a positive local transfer entropy.

Because we are interested in the time window where direct synaptical influences exist, we selected SLTE value at the time delay *d* when TE shows the peak value. We simply call the peak value of SLTE as *E-I bias* in the following sections. The *E-I bias* will hold the positive/negative sign coming from equation 3. Now, we can discriminate excitation or inhibition bias for individual connections in the two dimensional space of *Strength* (of TE) and *E-I bias* (of SLTE) (fig.3-(a), upper left panel). In order to define cell labels of excitatory or inhibitory *neurons* from signs given for individual *connections*, we plotted the average of the 1st-5th principal components and the average of the 6th-10th principal components of the *E-I bias* value among all output connections for individual all neurons in a two dimensional space, and categorized them into excitatory and inhibitory neurons by identifying their associated clusters. The averaging among several components helped to improve the signal-to-noise ratio, but the general trend was able to be captured even if we simply select the 1st and 2nd principal components, and the classification based on the axis of a single dimension of the sum of 1-10 components also provides almost the same result as the current two dimensional case. In the two dimensional space, especially, the cluster for excitatory neurons is prominent, so we selected the decision boundary which separates inhibitory neurons from excitatory neurons as a 2.5σ line from the center of the cluster for excitatory neurons.

**Figure 3.**
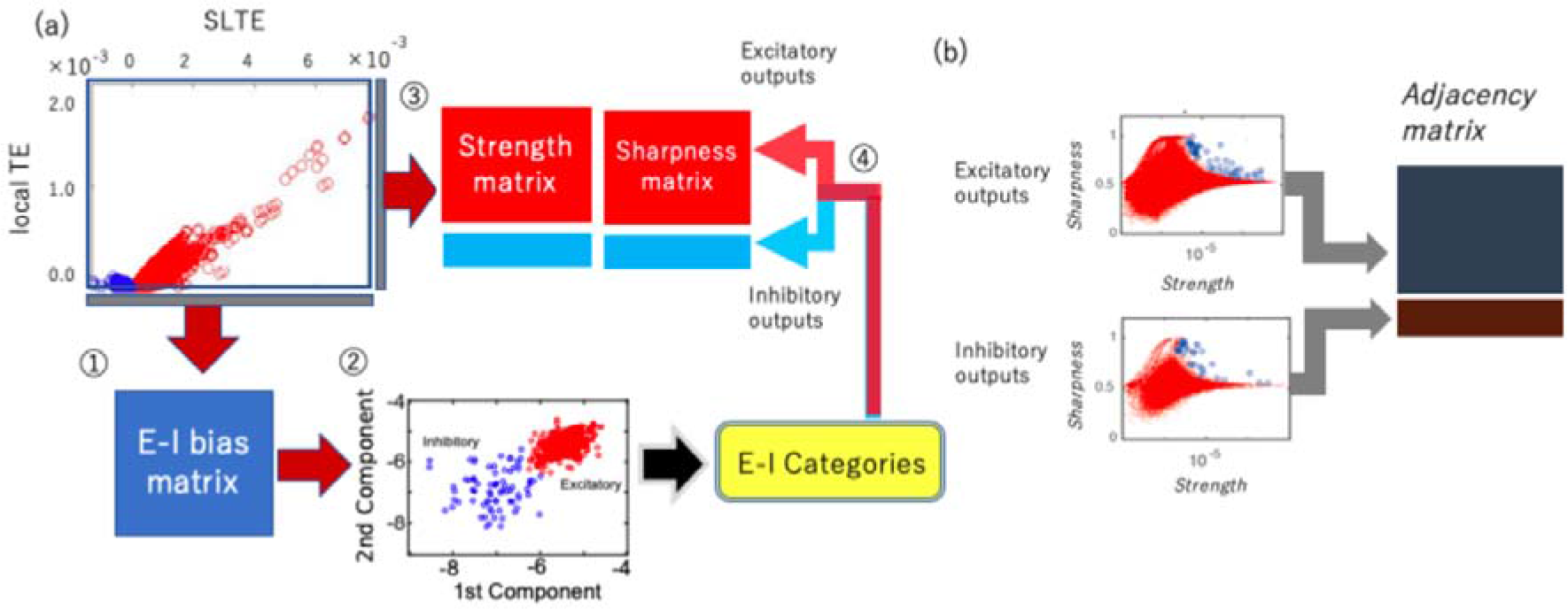
Combinational process of cell categorization and connectivity estimation: (a) and connectivity estimation (b): (a) The processing steps of cell categorization into excitatory and inhibitory neurons are listed as numbers here: ⍰ First, we calculated E-I bias (SLTE at the delay where TE shows the peak value). ⍰ Then, we determined the sign, positive or negative by observing the clustered distributions in the two dimensional space of the 1st and the 2nd SLTEs ??principal components??. ⍰ We calculated Strength and Sharpness from the TE histogram according to the analysis scheme shown in figure 2-(c,d). ⍰ Finally, we separated the Strength and Sharpness matrices into excitatory and inhibitory groups based on the classification of step ⍰. (b) The cell categorization preceded the connectivity determination because we have to prepare the shuffled null data separately for excitatory and inhibitory synaptic connections to determine connected neuron pairs [Refer to Shimono, Beggs, 2014 for more detailed information]. We separately produced the connectivity matrix for excitatory and inhibitory neurons by selecting strong or sharp connections as compared with the null shuffled data.

### 2-11. Defining sign and connectivity as a combination

The methods described in previous subsections 2-9 and 2-10 enabled us to get inhibitory and excitatory labels from the *E-I bias* matrix and the connection patterns from the *Strength* and *Sharpness* quantities. Now, we need to ask how we can systematically combine these two procedures, the Connectivity estimation (strengths, connected or not) and sign estimation (positive or negative)? The key point is that the cell categorization is necessary to be performed before we determine the connectivity, because we have to separately compare real data with the shuffled null data for excitatory and inhibitory synaptic connections. So, the starting steps ⍰ and ⍰ in Figure 3-(a) are purely taken for cell categorization, and the given cell categories are applied to the *Strength*, *Sharpness* matrices in the following step ⍰. Finally, as mentioned in subsection 2-9, we selected connections holding a strong and sharp peak in the TE histogram on the *Strength-Sharpness* space separately for excitatory and inhibitory connections (fig. 3-(b)). We generated 100 shuffled data by jittering spikes uniformly within a 19ms time window around the original time step when spikes existed. Here, notice that the sizes of the shuffling windows 19ms are the same among excitatory and inhibitory neuron groups. For the detailed individual procedures of cell categorization and the connectivity estimation, refer to the previous subsections 2-10 and 2-9 respectively.

### 2-12. Graph theoretical analyses

The estimation of connectivity matrix is not the final destination. The estimation was prepared to effectively untangle the complex web of interactions among huge numbers of neurons. This study specifically tries to understand the mechanism of how inhibitory neurons are able to maintain a relatively stronger influence in the microconnectome than excitatory neurons even though relative numbers of inhibitory neurons are much smaller than excitatory neurons. So, we should quantify the relative importance of individual neurons based on their functional connectivity pattern. We regard individual neurons as nodes, which equally include both excitatory and inhibitory neurons, and also regard a bundle of synaptic connections connecting a pair of nodes as links. Notice that in our study we could define directions for all individual links. Let us explain the basic network properties based on these components. The first step and the most naive evaluation of networks is degree or connectivity strengths, and the following second and third steps are quantifications of influences of individual nodes based on centrality and controllability respectively.

### 2-12-1. Degree

The degree tells how many connections extend around individual nodes. Degree can be categorized into in-degree and out-degree, which respectively express the numbers of input and output links directly connecting with individual nodes. So, the expression of in-degree and out-degree for a connectivity matrix are total number of input connections and total number of output connections by regarding matrix as a binary matrix. Histograms of input/output degrees are the key factor to know if there are hubs that have clearly more connections than other nodes [fig. 4-(a)].

**Figure 4.**
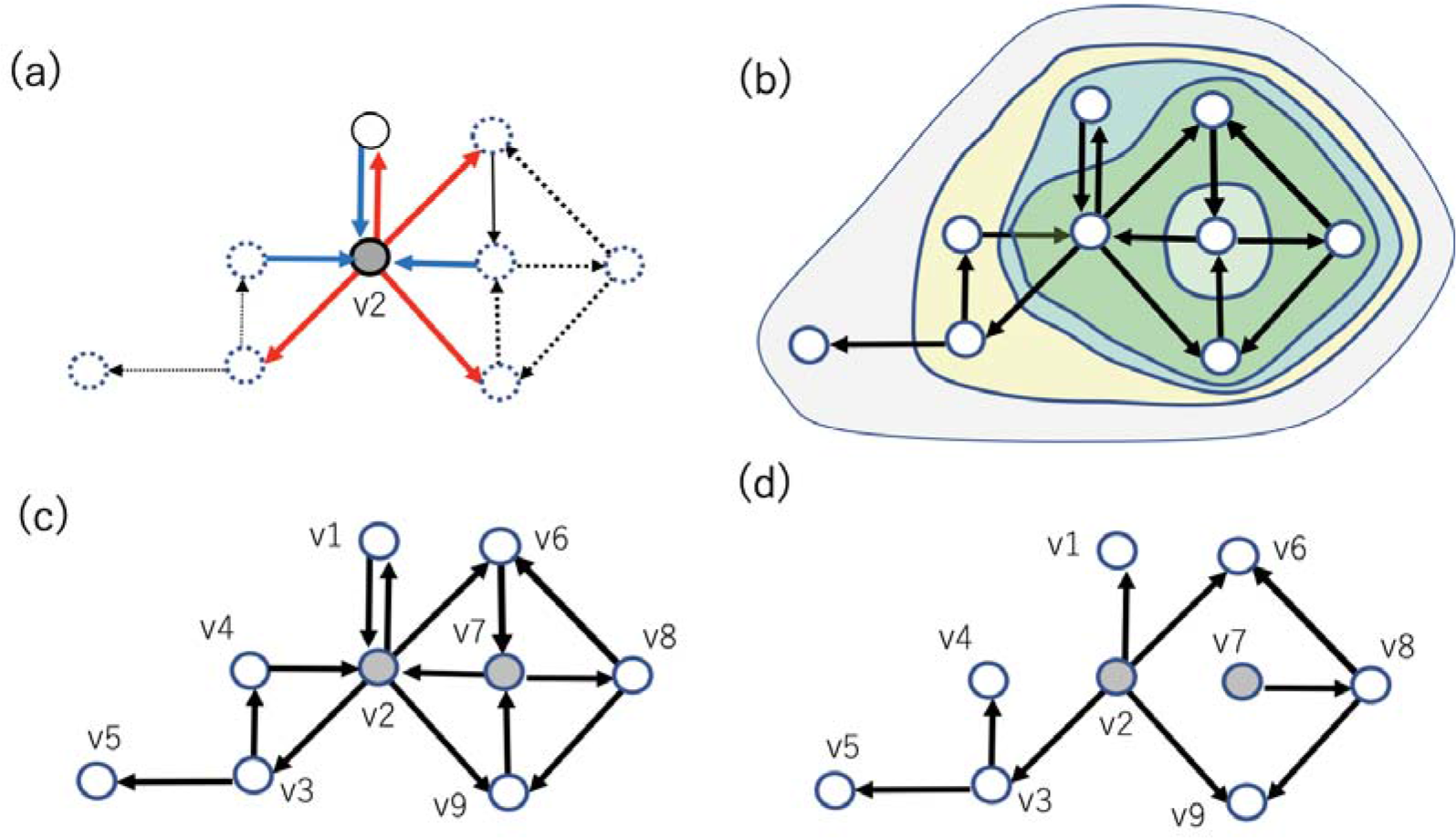
A schematic figure showing an example of network variables in a directed connectivity: (a) Out-degree or in-degree expresses how many links connect from or to a specific node respectively, shown as thick lines surrounding node v2. (b) An example of subsets of nodes extracted with a K-Core centrality algorithm. The stepwisely extracted subsets of nodes were surrounded by relatively smaller circles. (c) Here, an example feedback vertex set (FVS), nodes v2 and v7, are marked with gray color. FVSs like this example are defined as a set of nodes after eliminating inputs to the set, the remaining connectivity holds only directed cycles around the FVS. (d) If we remove incoming links to FVSs, all cycles included in the connectivity pattern disappear. This property represents that v2 and v7 are an example of FVSs in the system. The minimum number of nodes ?that are? FVSs in this example is 2. Therefore, v2 and v7 are also an example of minimum feedback vertex set (MFVS).

### 2-12-2. Evaluation of centralities with K-Core centrality

Broadly speaking, degree can be categorized as a kind of centrality measure, and the highly connected and well centralized specificity of hubs should be characterized based on many centrality measures. Here, we used two representative centrality measures. The first measure is K-Core centrality, which extracts the core, so subset of nodes, of more highly connected nodes than a threshold degree, and the evaluation of degree is stepwisely performed after peeling off lower degree nodes than the (k−1) [fig. 4-(b)]. Highly connected and highly central nodes will be simply called hubs.

### 2-12-3. Evaluation of Controllability via Feedback Vertex Set (FVS)

This subsection introduces our method for analyzing the architecture of effective networks from a viewpoint of *controllability*, more accurately speaking *structural controllability*. Here, controllability expresses the ability that we are able to guide a system from any initial state to any desired state in a finite time interval by stimulating *driver nodes*, which are nodes to whom external control inputs are directly given. Specifically, structural controllability focuses on the controllability of a given system characterized only with information of its static (or structural) architecture [Lin, 1974]; This approach held superiority to discussing whether a system is controllable for almost all parameters without knowing all parameters beyond the common problem that traditional control theory has often failed to identify parameters (e.g., coefficients) because of the property that traditional theories have utilized differential (or difference) equations to describe dynamics of the system.

Liu et al. (2011) recently introduced the concept of *structural controllability* into the research field of complex networks [Liu, et al., 2001], and many relating questions have been asked in applications of *structural controllability* for network systems; One key question there is how to select the minimum number of *driver nodes* to structurally control the whole system because it is realistically difficult that we simultaneously control a huge number of nodes by direct inputs to all of them.

In many cases this selection problem could be reduced to a graph theoretical problem, represented by approaches based on three characteristics: (i) maximum matching (MM) [Liu, et al., 2001], (ii) minimum dominating set (MDS) [Nacher, T. Akutsu, 2017], and (iii) feedback vertex set (FVS) [Akutsu, et al., 1998; Mochizuki et al., 2013]. Each model has some limitations. For example, controlling the targets for MM is limited to linear systems.

MDS usually requires several or more high degree nodes in order to control the system using only a small number of driver nodes.

The target (desired) states of FVS should be (statically or periodically) stable. The hidden dynamics of neurons behind the connectivity matrix show strong non-linearity, and also do not have many <u>high degree nodes</u> because long-tailed distribution could be observed. So, we adopted the FVS-based approach here.

Now, let’s see the example shown in figure 4-(c), and imagine the situation when we cut all links incoming to nodes v2 and v7 from the original connectivity pattern (fig. 4-d). Then, you will realize that the remaining connectivity will lose all *directed cycles*, *which* are closed loops or cycles created by connecting directed links in series (Fig. 4-d). Such a subset of nodes like the set of {v2,v7} are called *feedback vertex sets* (FVSs).

About the situation of figure 4-f, you will also realize that all other nodes are located “downstream” of FVSs (like {v2,v7}). This relative position of FVSs says that, when we control the value of FVSs as constant, the dynamics of the other nodes will also become sequentially entrained into one stable state after passing a long enough time interval. This property suggests that we can use FVS as a *driver node set* if the desired state is a steady state. Even for the case of figure 4-(d) when the cut links still remain, we will be able to control the state into a set of periodic states. So, FVSs can play the role of a *driver node set* even when the desired state is a set of attractors.

Remember that the current target is to seek the minimum combination of *driver nodes*. So, we specifically want to seek the smallest FVSs, consisting of the minimum number of nodes, which is called *minimum feedback vertex sets* (MFVSs). Because we would generally not try to drive a system to non-steady states, it is a realistic demand that we seek MFVS, as a minimum size driver node set. The FVS-based approach can be also applied to wider classes of non-linear systems as well as linear systems [Mochizuki et al., 2013]. Furthermore, the FVS-based approach was applied to the analyses of gene regulatory networks [Mochizuki et al., 2013; Zanude et al. 2017] and signaling pathways [Bao et al., 2018], and the results suggest that nodes in MFVS or nearly minimum feedback sets often correspond to genes or proteins playing important biological roles.

Here, we should notice that MFVS is not necessarily unique; For example, the set of {v2,v8} is also a MFVS. So, we would be better to categorize nodes that are part of MFVSs into two groups, one group is nodes participating in all MFVSs [Bao et al., 2018], and another group is nodes participating in one of the MFVSs but not in all of them. These are called *critical and intermittent* nodes respectively, and the remaining nodes are called *redundant* nodes [Jia et al., 2013]. In the example shown in figure 4-d, v2 is the only one *critical* node by chance, v7, v8, v9 are *intermittent* nodes, and others are *redundant* nodes.

Currently, it is known that the computation of a MFVS is NP-hard, which implies that it is not plausible that there exist theoretically efficient algorithms for finding a MFVS. However, it is possible in practice to compute an MFVS for up to moderate size networks using Integer Linear Programming (ILP). We employed this simple ILP formalization to compute MFVSs and types of nodes, using IBM ILOG CPLEX as a solver for ILP.

## 3. Results

### 3-1. Basic properties

First, we observed the histograms of degrees, number of connections, of individual neurons for the excitatory and inhibitory cases. Figure 5-a shows the histogram on a log-log space. The result shows that the degree histogram follows a long-tailed form and that there are hubs, nodes having exponentially more numbers of connections than others, not only for excitatory neurons but also for inhibitory neurons [fig. 5-a]. The existence of hubs for excitatory effective connectivity had been already reported in our past studies [Shimono, Beggs, 2014].

**Figure 5.**
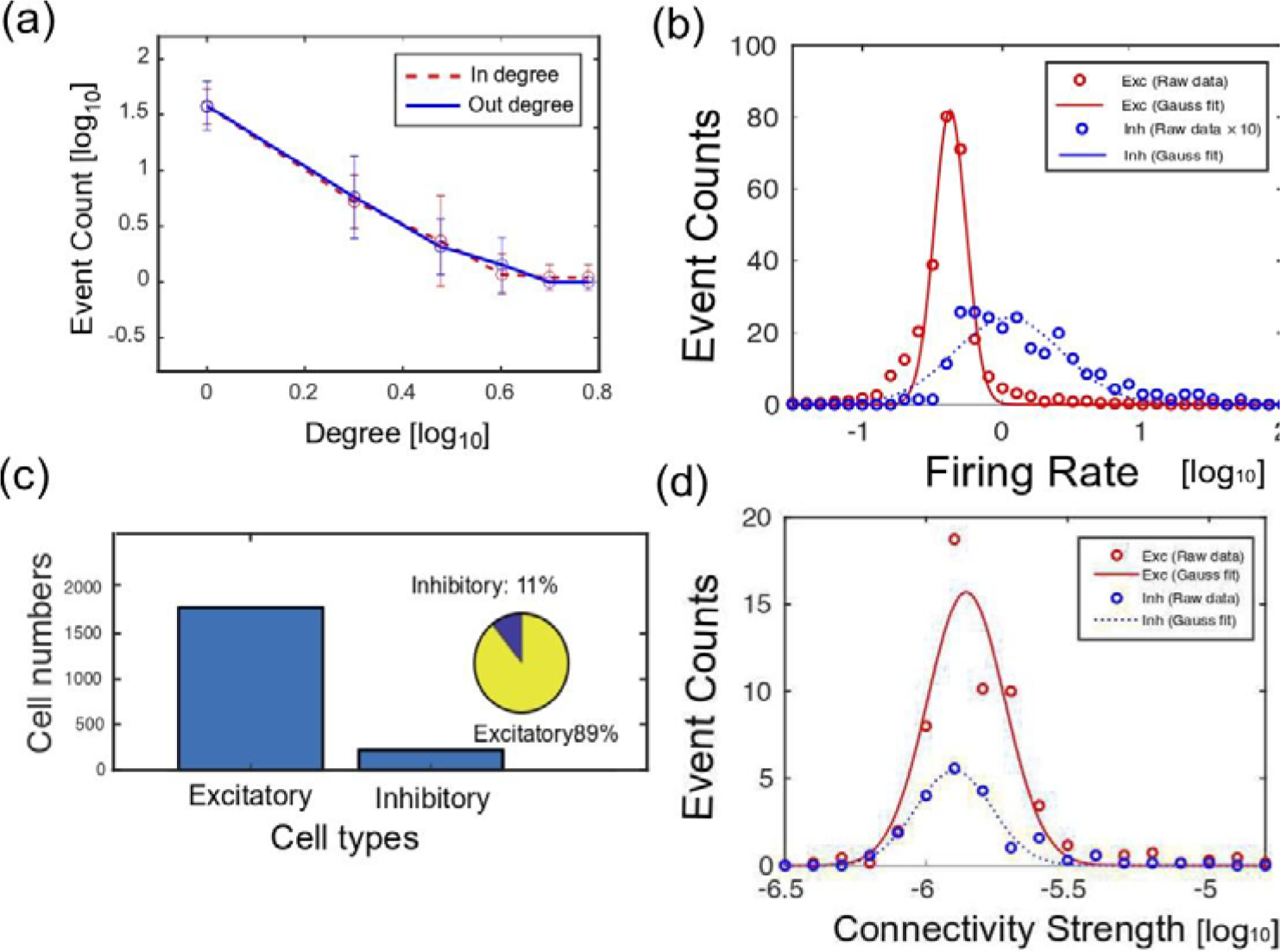
Basic properties of neuronal networks: (a) shows the degree histogram, so histogram of number of connections. The solid line and dotted line show results for out degree and in degree for all neurons respectively. (b) shows histograms of firing rate for excitatory neurons (solid line) and inhibitory neurons (dotted line). (c) The main bar graph shows the total number of identified excitatory and inhibitory neurons. The inserted pie chart shows the relative ratio of their numbers. (d) shows histograms of connection strengths for excitatory neurons (solid line) and inhibitory neurons (dotted line). The x-axis of panels (a), (b) and (d) are log scale with base 10.

Second, we started to observe the rules the activities of excitatory and inhibitory neurons follow. Their firing rates approximately followed log-normal distributions. Although the log-normal distribution of the firing rate among excitatory neurons is a relatively well-known basic property, the log-normal rule of firing rate among inhibitory neurons is not well-established yet. As a similar aspect in terms of log-normal rule, we observed the distributions of connectivity strengths of effective networks [Nigam et al., 2014]. This study revealed that the connectivity strengths not only among excitatory connections but also among inhibitory connections follow a log-normal rule [fig. 5-d]. The relative ratio of numbers of excitatory neurons was around 90% [fig.5-c].

### 3-2. Estimations of influential neurons based on centrality [k]

Because we could find exceptionally high degree nodes in the system, we wanted to evaluate the influence of individual nodes. The first step to quantify their influence is to evaluate how central the position of the individual neurons are. To characterize such centrality of individual neurons, we adopted K-core centrality here [Fig. 6]. Then, we could observe that the averaged values of K-core centrality of inhibitory neurons are higher than ones of excitatory neurons for all data samples for all 7 cortical slices [Fig. 6-a]. The superiority of inhibitory neurons in terms of centrality could be especially observed in deeper layers, so layers 4-6 [Fig. 6-b].

In summary, we found that influential neurons in terms of centrality are inhibitory cells, and that such highly influential inhibitory neurons are mainly observed in deep layers, 4-6 layers, and especially layer 6 showed a significant difference.

**Fig. 6.**
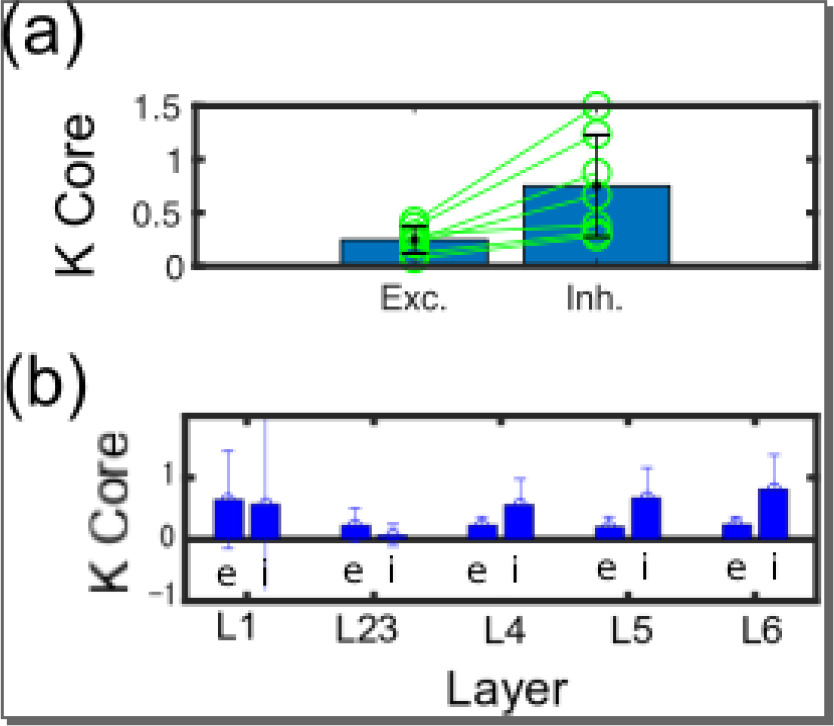
E/I cell category and K core centralities: (a) shows the difference of averaged K core values for all excitatory and inhibitory neurons. The error bars and circles correspond with the variety among 7 cortical slices. They showed significant difference (Wilcoxon paired test; p<0.05). (b) shows the difference of averaged K core values for excitatory and inhibitory neurons within individual layers. The left and right bars for individual layers correspond with excitatory and inhibitory neurons respectively.

### 3-3. Estimations of influential neurons based on FVS

We also quantified the influence of neurons based on their ability to control other neurons based on FVS. The main result is shown in figure 7. We could find the relative influence of inhibitory neurons as significantly stronger than excitatory neurons (p<0.05; Wilcoxon paired test). Additionally, we also observed the FVS values between excitatory and inhibitory neurons separately among different layers. Then, we could find the superiority of inhibitory neurons at deep layers (layers 4-6) and especially could find a significant difference at layer 6.

Finally, we compared the relative commonality between FVS neurons and neurons holding high K Core centralities, and could find the significant statistical inter-dependency between these two types of influential neuron pools.

**Fig. 7.**
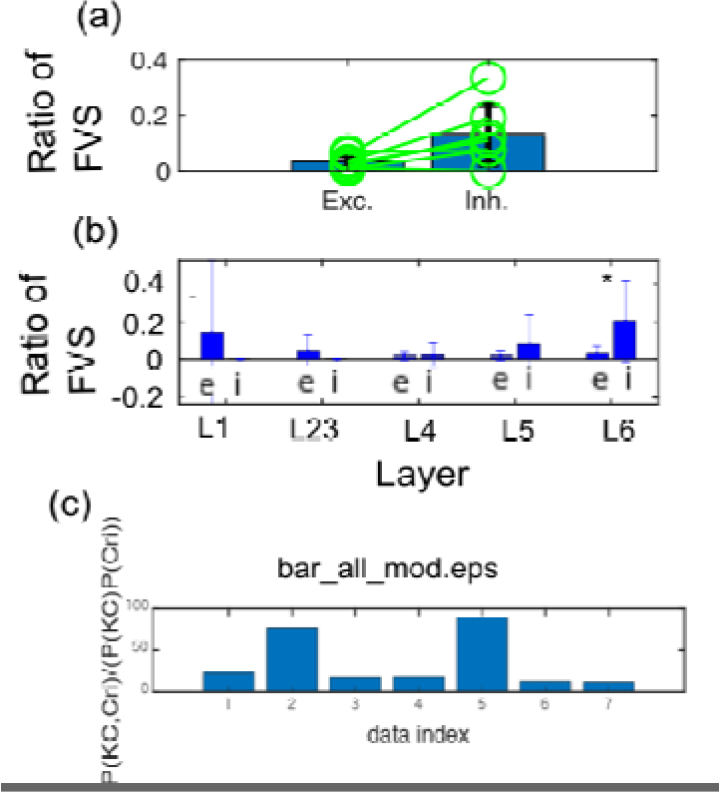
E/I cell category and FVS: (a) shows the difference of Ratio of number of FVS within excitatory or inhibitory neuron pools in the same manner as figure 6-(a). The E/I difference also showed significant difference (Wilcoxon paired test; p<0.05). (b) shows that while Layers 1 to 5 didn’t have any significant difference in the ratio of FVSs among excitatory and inhibitory neurons, we could find significantly more inhibitory FVSs in Layer 6.

## 4. Discussion

### 4-1. Brief summary

The brain is a highly non-uniform network system. We sought to answer how inhibitory neurons are able to keep the balance against excitatory neurons, which are more abundant than inhibitory neurons by seeing the neuronal network as a non-uniform network system. This study has tried to move forward to reveal how the stronger influence of inhibitory neurons is supported within the network topology of functional interactions among ~1000 neurons beyond simple differences such as among firing rates.

### 4-2. Log-normal rules

First, we started to observe the difference of firing rate and connectivity strengths of effective connectivity between excitatory and inhibitory neurons. These parameters were previously observed only for excitatory neurons [Shimono, Beggs, 2014; Nigam et al., 2015]. In this study we tried to add more information by separately observing excitatory and inhibitory neurons. We could observe that the firing rate in inhibitory neurons is higher than excitatory neurons, as known before. Additionally, we could commonly observe log-normal like distributions of firing rate and connectivity strengths for inhibitory neurons. Past computational simulation study demonstrated that a Hebbian learning rule can produce the situation where both firing rate and connectivity intensity show log-normal distributions [Koulakov et al., 2009]. However, the experimental tests for inhibitory neurons have been few. A simple mechanism causing the phenomenon that effective connectivities obey log-normal distribution might be that the underlying “structural” connectivity strengths, reflecting the spine sizes, obey a multiplicative rule [Loewenstein et al., 2011]. However, more study will be still necessary to clarify relationships between spine sizes and strengths of effective connectivity in the future.

In summary, the finding that inhibitory neurons obey log-normal rule hasn’t been well-established. Our finding is new and provides several suggestions. But, we started to ask more questions about the relative positions of E/I neurons in the network topology.

### 4-3. Influence of neurons in terms of Centrality and Controlling ability

When we observe the histograms of link numbers of individual neurons, so degree histograms, we see that they are long-tailed. From this point, past study demonstrated that there are hubs among excitatory neurons [Roxin et al., 2008; Shimono, Beggs, 2014], and rich club organization [Nigam et al., 2016]. This study additionally demonstrated that there are hubs even if we include inhibitory neurons in the effective networks. From the basic observations, we moved the main focus on the difference of relative positions of inhibitory and excitatory neurons in the effective networks, and we quantified the relative difference of influence of inhibitory and excitatory neurons according to how central these neurons are located. We could observe that inhibitory neurons take more central positions than excitatory neurons in terms of K-Core centrality. Additionally, we could find that the significantly more central property of inhibitory neurons than excitatory neurons could be observed in deep cortical layers by combinational evaluations of the effective networks with NeuN immunochemistry staining.

Besides, we also tried to extract the most “influential” neurons in terms of their controlling ability of other neurons by extracting the minimum combination of Feedback Vertecs. Then, we could consistently observe the stronger controlling ability of inhibitory neurons compared to excitatory neurons, and interestingly, again the influential inhibitory neurons are selectively located in deep cortical layers. This result also suggests that, if we want to effectively regulate the behavior of a neuronal system, stimulation of inhibitory neurons might be more influential than excitatory neurons.

In summary, we could reveal that inhibitory neurons not only are located at more central positions than excitatory neurons, but also have higher controlling ability of the entire network dynamics in local cortical neuronal circuits, in terms of FVSs. So, inhibitory neurons play a role of a central controlling regulator.

### 4-4. Final remarks

The Brain consists of a huge number of components, neurons, and their connections are intertwined with each other in a very complex manner. In order to understand the system properties, we need to unwind the complex connectivity. This study asked how inhibitory neurons play different roles than excitatory neurons and keep balance with them, and we find the specific characteristics of inhibitory neurons in terms of centrality and controlling ability of other neurons. We also find that such highly central and controller inhibitory neurons are selectively located at deep layers. This statement is very simple, but it was achieved by utilizing cutting-edge recording technology, and sophisticated accumulations of analysis methods. This finding and research approach will develop many related studies on various cell categories, brain regions, disease states, animal species and so on, and also provide good constraints to develop realistic computational models in the near future.

## Acknowledgements

MS is supported by several MEXT fundings (19H05215, 17K19456) and Leading Initiative for Excellent Young Researchers (LEADER) program, and grants from the Uehara Memorial Foundation. The MRI experiments of this work were performed in the Division for Small Animal MRI, Medical Research Support Center, Graduate School of Medicine, Kyoto University, Japan. We warmly acknowledge for all supports by Hakubi center, Marie Obien, Urs Frey, Hirohiko Imai, and Pierre Yger to establish this study.

## Notes

**Conflict of interest** The authors declare no competing financial interests

